# Identification of Spinster Homolog 3 as a novel putative LL-37 receptor

**DOI:** 10.1101/2023.07.07.548096

**Authors:** Alana Bazán Corrêa, Thais Martins de Lima, Suely Kubo Ariga, Fabiano Pinheiro da Silva

## Abstract

**Background:** Antimicrobial Peptides are primitive components of the innate immune response. LL-37, a human antimicrobial peptide, has been researched for its therapeutic potential in immune-inflammatory diseases. This study aimed to identify novel LL-37 receptors.

**Methods and Results:** Using the phage display technique, the sequence GNWSFV was selected and found to bind to LL-37. BLAST analysis revealed that this sequence has a high degree of similarity to a portion of human transmembrane protein Spinster Homolog 3.

**Conclusion:** Our study suggests that Spinster Homolog 3 may be a potential therapeutic target for LL-37-related diseases.

## Introduction

Antimicrobial peptides (AMPs) are a large group of compounds produced by multicellular organisms in both the animal and plant kingdoms. These peptides act in host defense and are a primitive and conserved component of the innate immune response, acting against a large number of microorganisms, including bacteria, fungi, yeasts and viruses [1].

Antimicrobial peptides vary in their sequence and structure. They are usually amphipathic, contain from 12 to 50 amino acids, and have a net positive charge. These chemical properties allow them to insert into the cell membranes of invading organisms, leading to their death. In addition to their antimicrobial properties, AMPs also have immunomodulatory effects, regulating the activities of various immune cells, including macrophages, neutrophils, T cells, B cells, and dendritic cells [1].

Cathelicidins are a family of AMPs with one single member in humans, named LL-37. It has been implicated in a range of diseases, including autoimmune diseases, cancer, and infectious diseases. It has been studied extensively for its potential therapeutic applications, including as a novel antimicrobial agent [2] and as an immunotherapy [3].

Even though its mechanisms of action remain poorly understood [4], it has been described that LL-37 can bind to several unrelated membrane receptors, such as N-formyl peptide receptors (FPR1/2) [5], CD11b/CD18 [6] and CXCR2 [7], among others.

The aim of the present study was to identify novel LL-37 receptors, through the phage display technique. Phage Display is a cloning technique that allows the expression and selection of small peptide sequences on the surface of bacteriophages [8]. It is a versatile technique that frequently reveals previously unknown molecular interactions, putting in evidence new signaling mechanisms and therapeutic targets.

## Materials and Methods

### Phage display

The phage display technique is based on the construction of bacteriophage libraries that contain small peptide sequences (between 5 and 15 amino acids) displayed on one of the viral capsid proteins [9].

Synthetic LL-37 was obtained commercially (95% or greater purity). The library contained an estimated diversity of 8 × 10^9^ distinct inserts, and the technique was performed as previously described by our group [10].

Briefly, library and individual phages were cultivated by infection with *E. coli* strain K91kan grown in Luria–Bertani (LB) medium supplemented with kanamycin (100 μg/mL) and tetracycline (20 μg/mL) and purified from the supernatant of the cultures using the PEG/NaCl method [11].

LL-37 was immobilized in 96-well plates (1 μg in 50 μL of PBS) overnight at 4°C. Wells were then washed with PBS and incubated with the phage library (10^9^ bacteria transducing units in 50 μL of PBS). After 2 h, the wells were washed with PBS and phage bound to the immobilized receptor were recovered by infection with *E. coli* K91kan at log phase and quantified by colony counting. Phages were amplified and used for the next round of selection. After four rounds of selection, phage clones were submitted to amplification and selected for DNA sequencing to determine the peptide encoded by each individual phage clone (**Figure 1**). In the end, the sequences were submitted to binding assays, in the presence of LL-37 or a scrambled peptide. The binding assays confirmed that the sequence GNWSFV binds to LL-37. Bovine serum albumin (BSA) and scrambled peptide were used as controls (**Figure 2**).

**Figure 1.**
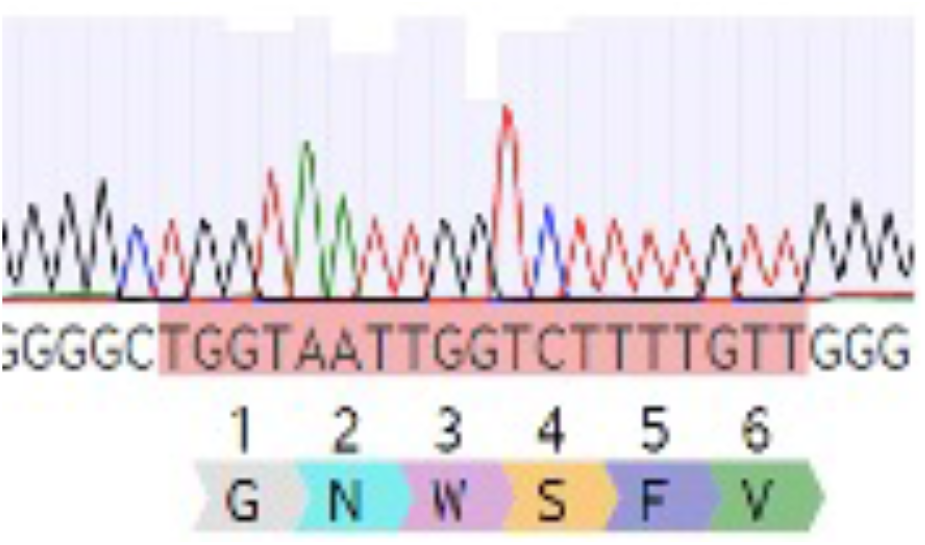
DNA sequencing of the clone selected via biopanning.

**Figure 2.**
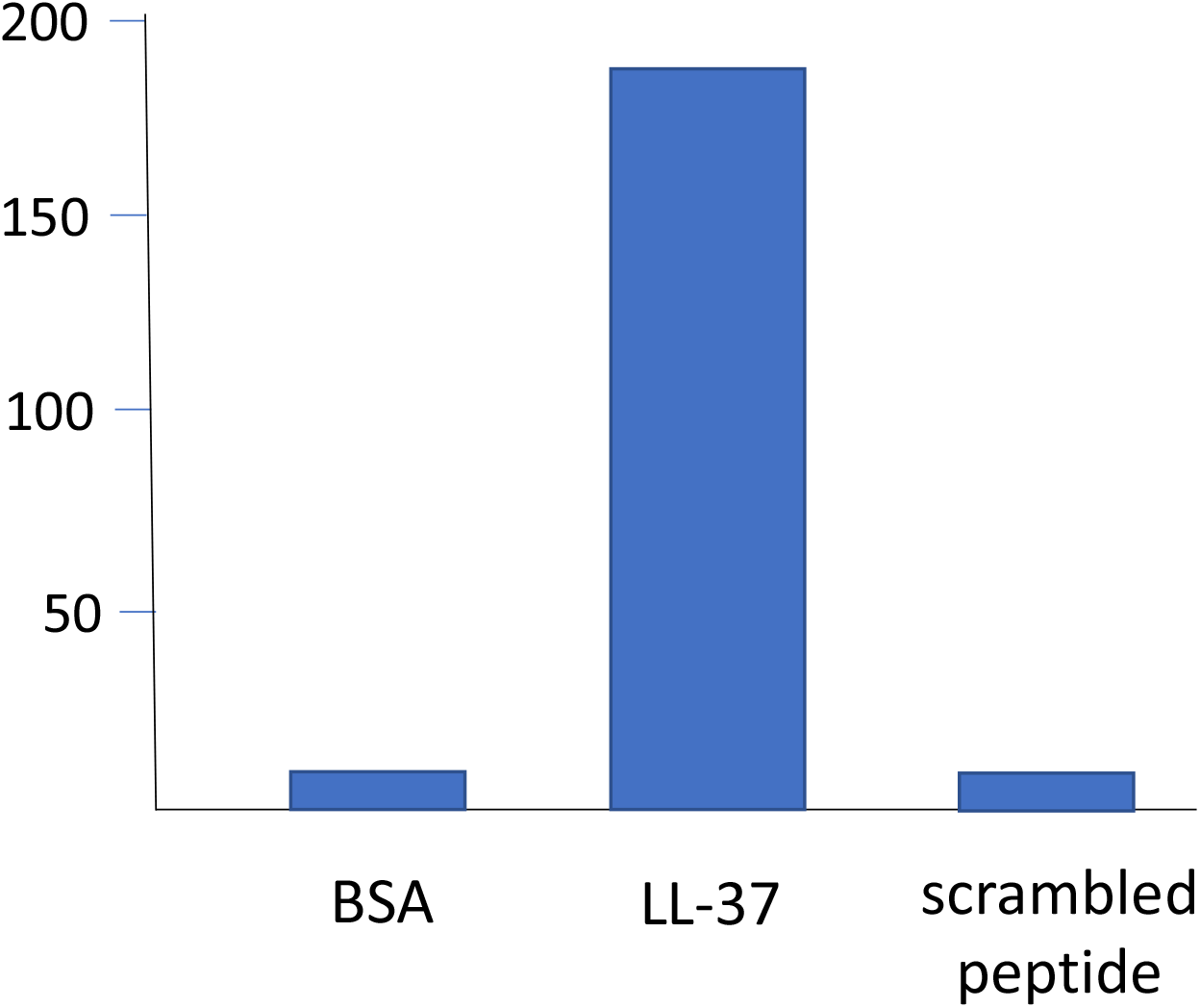
Binding assay confirming that the sequence GNWSFV binds to LL-37.

### BLAST

The sequence GNWSFV was analyzed by BLAST.

BLAST is a tool that finds regions of similarity between biological sequences, thus comparing nucleotide or protein sequences with sequence databases (https://blast.ncbi.nlm.nih.gov/Blast.cgi).

## Results and Discussion

BLAST search detected that human Spinster homolog 3 (SPNS3), a transmembrane transporter, contains a protein sequence with the highest score of similarity and the lowest E value, when compared with the sequence GNWSFV. The twenty highest scores are shown in **Table 1**.

**Table 1.**
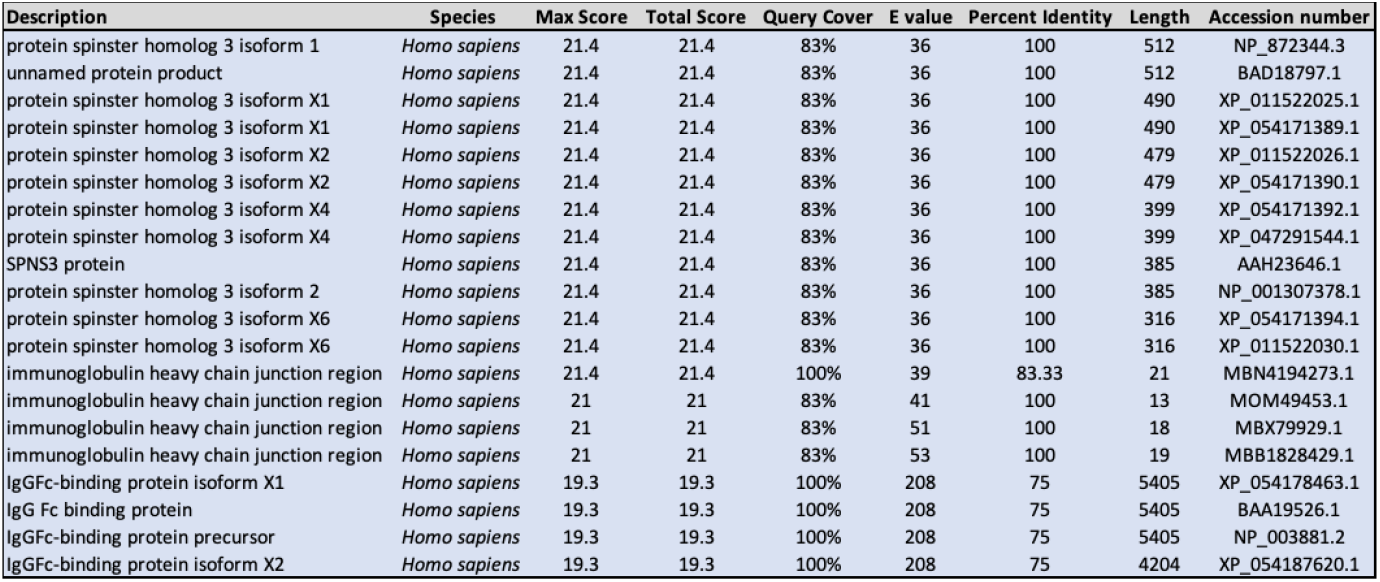
List of the top twenty proteins that exhibit sequences with the highest score of similarity and the lowest E values, when compared to the amino acids sequence GNWSFV. Results obtained via a BLAST search.

Human spinster homologue 3 is a widely distributed transmembrane protein predicted to be involved in the transport of sphingolipids. It was identified in 2007 due to its structural similarity with the solute carrier family 22 (SLC22), a large family of cation and anion transporters of the major facilitator superfamily [12]. Few studies investigated this protein, and its role in human diseases remains unclear, but there is some evidence that it is implicated in cell death mechanisms [13], through the autophagy-lysosome pathway [14, 15]. Indeed, SPNS2 and SPNS3 have been appointed as biomarkers of apoptosis resistance and poor prognosis in acute myeloid leukemia (AML) [16, 17].

Studies suggest that human spinster homologue 3 may participate in the sphingosine-1-phosphate signaling pathway (S1P), transporting S1P to the extracellular milieu to activate downstream signaling [16, 17]. S1P mediates several cellular processes, such as proliferation, survival, migration, angiogenesis, among others. Interestingly, S1P stimulates LL-37 gene expression and protein release [18].

Cholesterol- and sphingolipid-enriched membrane domains exist either as “lipid rafts” or, when associated with caveolin, as invaginations called caveolae. It has been shown that LL-37 interacts with these domains [19]. We suggest a new signaling mechanism, where SPNS3 induces LL-37 signaling via the S1P pathway. LL-37, then, binds to SPNS3 directly, to fine tune its activation. Since LL-37 has both activating and inhibitory effects [20], further studies are necessary to answer if this a positive or a negative loop.

## Conclusion

Our results showed that the sequence GNWSFV binds to LL-37 and was found to have a high score of similarity with a portion of the human transmembrane protein Spinster Homolog 3 (SPNS3). SPNS3 is involved in the transport of sphingolipids and has been implicated in cell death mechanisms, as well as in the sphingosine-1-phosphate (S1) signaling pathway. Our study suggests that LL-37 participates in SPNS3 signaling through direct binding and that SPNS3 may be a potential therapeutic target for diseases where LL-37 plays a role.

## Statements & Declarations

### Author contributions

FPS conceived and supervised the study. The experiments were performed by ABC. TML and SKA provided technical support. The first draft of the manuscript was written by ABC. FPS wrote the final version.

### Ethics approval

Protocols were in accordance with the University of São Paulo Faculty of Medicine Ethical Committee (project number 1179/2018).

### Data availability statement

All data generated during this study are available on reasonable request.

## Acknowledgements

We would like to express our gratitude to Prof Ricardo José Giordano for sharing his phage display library and for the technical guidance.

